# Energy filtering enables macromolecular MicroED data at sub-atomic resolution

**DOI:** 10.1101/2024.08.29.610380

**Authors:** Max T.B. Clabbers, Johan Hattne, Michael W. Martynowycz, Tamir Gonen

## Abstract

High resolution information is important for accurate structure modelling. However, this level of detail is typically difficult to attain in macromolecular crystallography because the diffracted intensities rapidly fade with increasing resolution. The problem cannot be circumvented by increasing the fluence as this leads to detrimental radiation damage. Previously, we demonstrated that high quality MicroED data can be obtained at low flux conditions using electron counting with direct electron detectors. The improved sensitivity and accuracy of these detectors essentially eliminate the read-out noise, such that the measurement of faint high-resolution reflections is limited by other sources of noise. Inelastic scattering is a major contributor of such noise, increasing background counts and broadening diffraction spots. Here, we demonstrate that a substantial improvement in signal-to-noise ratio can be achieved using an energy filter to largely remove the inelastically scattered electrons. This strategy resulted in sub-atomic resolution MicroED data from proteinase K crystals, enabling accurate structure modelling and the visualization of detailed features. Interestingly, filtering out the noise revealed diffuse scattering phenomena that can hold additional structural information. Our findings suggest that combining energy filtering and electron counting can provide more accurate measurements at higher resolution, providing better insights into protein function and facilitating more precise model refinement.

---

At atomic resolution, commonly defined as beyond 1.2 Å, individual atoms are fully resolved revealing an accurate model of the underlying structure^1^. Unfortunately, noise in the measurement often makes it difficult to obtain such high-resolution information from macromolecular crystals. The mean diffracted intensity decreases rapidly at higher resolution, and radiation damage limits the total information that can be recovered from each crystal. This necessitates the use of a low fluence. The signal-to-noise ratio is therefore poor, limiting the attainable resolution. Sufficiently strong data can still be retrieved in microcrystal electron diffraction (MicroED)^2^, supported by the use of fast and highly sensitive cameras that are effective even at low flux densities^2–4^. Recently, we reported a substantial improvement in data quality by recording data using electron counting on direct electron detectors^5^. The improved accuracy and resolution enabled *ab initio* phasing using a fragment-based approach^5^, and allowed identification of hydrogen bond networks and protonation state of atoms^6^. However, this was demonstrated for triclinic lysozyme, which is relatively small and forms crystals with low solvent content, while proteinase K, which is much larger and contains more solvent, did not reach similarly high resolution. Electron counting detectors do not eliminate all noise, and further optimization of the experimental setup and data collection strategies is needed to routinely obtain more high-resolution information.

Inelastic scattering is a major factor limiting the signal-to-noise ratio^7,8^. Upon an inelastic interaction, electrons lose a small fraction of their energy and are no longer coherent. This contributes to increased background noise and a broadening of the Bragg peaks, thereby affecting the kinematic diffraction signal. This poses a significant challenge, as inelastic events are 3-4 times more likely than elastic scattering in biological specimens^8,9^. Whereas lattice filtering or subtracting a smooth radial background can partially correct for this^10,11^, post-processing does not reduce the noise inherently present in the measurement and cannot recover weak high-resolution reflections that are at or beneath the noise level. Removing inelastically scattered electrons experimentally based on their energy loss is therefore preferable. Previous reports assessed the effects of energy filtering on macromolecular diffraction data, demonstrating a significant reduction in background noise and sharper spots^3,12,13^. Interestingly, whereas the intensity and model statistics of the filtered data improved, the resolution of the reported structure did not^13^.

We hypothesized that combining energy filtering with the improved accuracy and sensitivity of direct electron detection would enhance data quality *and* resolution in MicroED. Two important considerations to this approach are the total fluence that can be withstood by the crystals, and the flux tolerated by the camera before coincidence loss prevents an accurate representation of the counts. The diffraction signal therefore is proportionally weaker. To mitigate this, we devised the following protocol that continuously rotates the crystals slower, uses the same total fluence spread over a smaller wedge, and utilizes the energy filter to further improve the signal-to-noise ratio. We applied this strategy to collect electron-counted and energy-filtered MicroED data from crystalline lamellae of proteinase K that were machined to an optimal thickness of ∼300 nm^14^. Previously, a maximum resolution of 1.4 Å was achieved from similar lamellae without the use of energy filtration^15^.

Initial diffraction images were taken to assess lamellae quality and achievable resolution (Fig. 1). Diffraction spots extended to the edge of the detector beyond 1.0 Å resolution, showing less background noise and sharper Bragg peaks compared to unfiltered data (Extended Data Fig. 1)^5,15^. Interestingly, energy filtering revealed diffuse scattering from the bulk solvent that is largely obscured in unfiltered data (Fig. 1)^5,15^. Continuous rotation MicroED data were collected from the best crystals, showing high quality information and spots extending out ∼1.0 Å resolution (Extended Data Fig. 2-3). Individual datasets were processed and showed significant information up to 1.06 Å resolution (Extended Data Fig. 4). Not all lamellae diffracted to the same resolution depending on crystallinity, quality of the lamellae and lattice orientation. Data from 17 crystals were merged and truncated at 1.09 Å resolution to ensure high multiplicity and complete sampling of reciprocal space (Extended Data Table 1). Intensity statistics show a clear improvement in attainable resolution and a more gradual decrease in crystallographic quality indicators compared to unfiltered data (Fig. 2A)^15^. Importantly, improving the resolution more than doubled the number of unique reflections used for structure refinement (Extended Data Table 1). At atomic resolution, the electrostatic potential map showed highly resolved features, enabling accurate interpretation of the structural model and visualization of hydrogen atoms (Fig. 2B).

**Figure 1.**
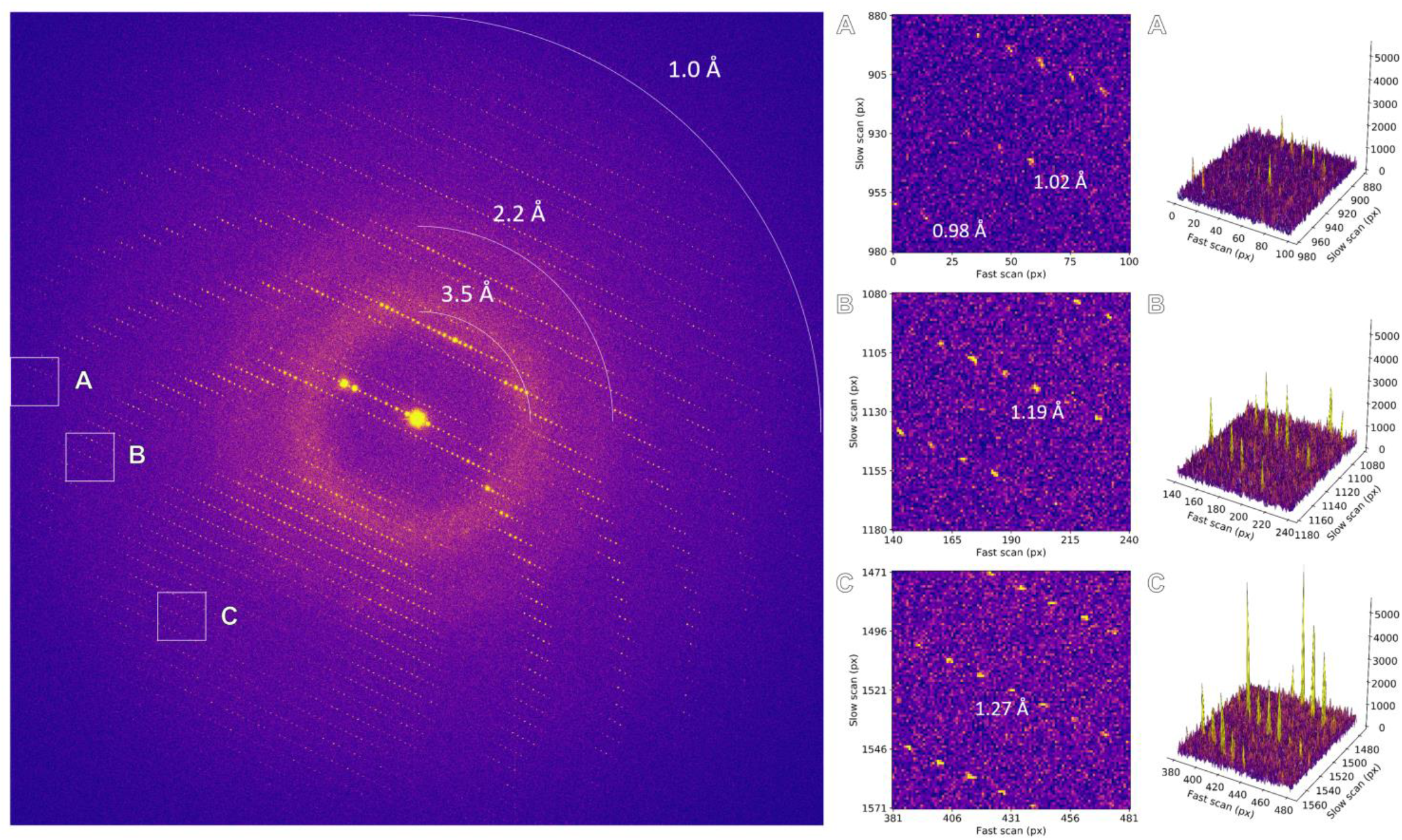
Energy-filtered diffraction pattern of a proteinase K lamella. Areas highlighted by the insets are magnified and the corresponding peak profiles are plotted.

**Figure 2.**
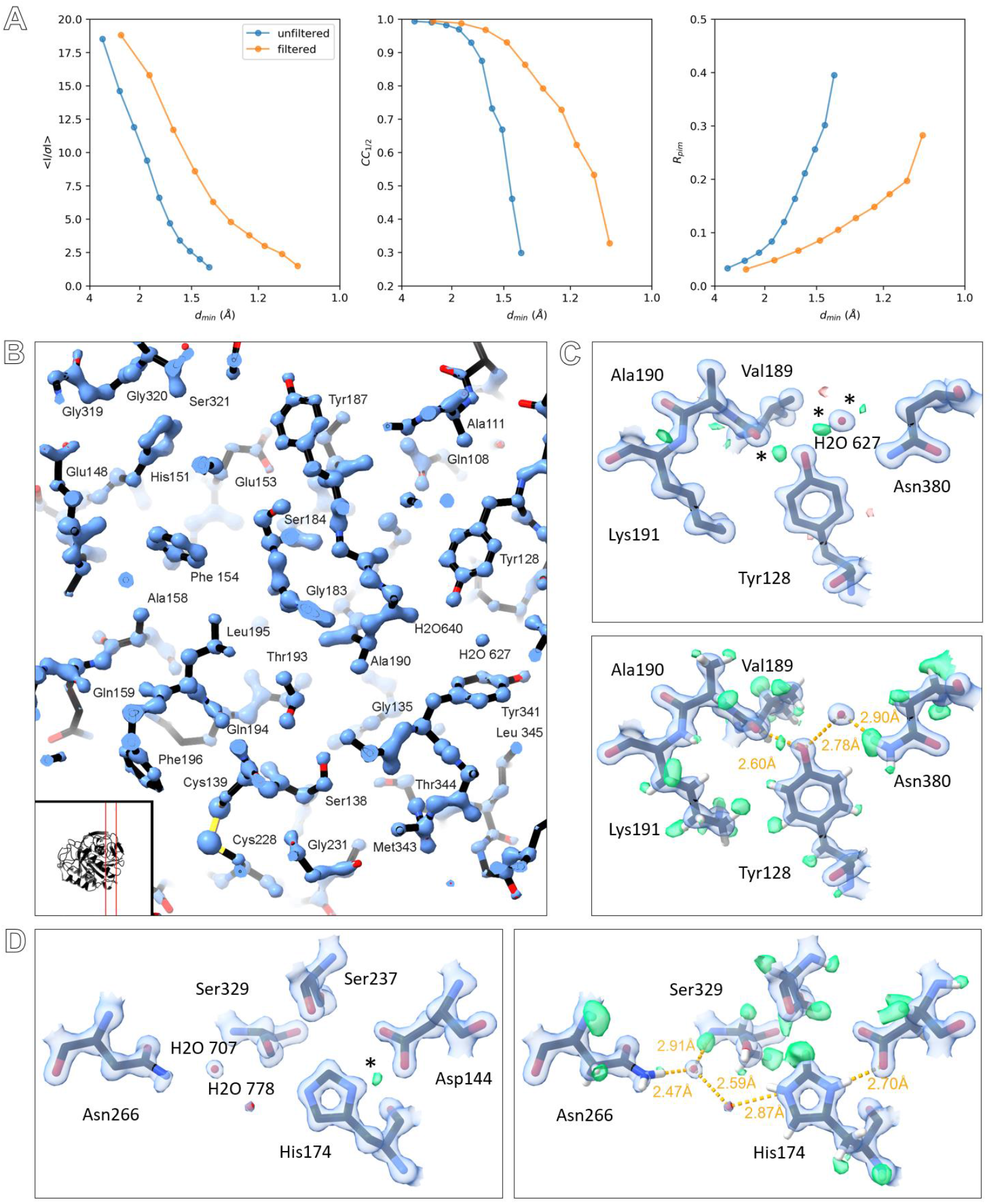
A. Comparison of intensity statics for filtered and unfiltered MicroED data, featuring from left to right: *I/σI*, CC_1/2_, and Rpim. B. Slice through the structural model showing the electrostatic potential map, the location of the slice in the structure is indicated in the inset. C. Maps and hydrogen omit maps shown for Tyr128. D. Maps and hydrogen omit maps for active site residues and waters. Electrostatic potential 2mFo-DFc maps are shown in blue and contoured at 4σ (B), and 2.2σ (C-D). Difference mFo-DFc maps are shown in green and red for positive and negative density contoured at 3σ (C-D). Hydrogen omit maps are shown in green and contoured at 2.3σ (C-D). Difference peaks marked with an asterisk indicate potential hydrogen atoms that have not been included in riding positions prior to map calculations.

We demonstrate that optimizing MicroED data collection by integrating an energy filter with direct electron detection cooperatively increases data quality and resolution by mitigating the effects of inelastic scattering. The filtered data exhibited reduced background noise, sharper spots, and an improved signal-to-noise ratio. Better high-resolution data provides more detailed insight into protein structure and function, and may prove useful in visualization of hydrogen atoms and hydrogen bonding networks^6^, as well as charge distribution^3,10^. With this improved data collection approach, the attainable resolution for proteinase K is similar to the best reported proteinase K structure by X-ray crystallography^16^, indicating that as technologies improve MicroED can rival more established structural biology methods. Diffraction spots extended beyond 1.0 Å resolution on individual frames, suggesting that there is still room for improvement. A data collection strategy that combines a larger number of datasets^3^, or a serial approach using individual snapshots^11^, may further improve data quality and resolution. Furthermore, the possibility to separate inelastic scattering from the signal is unique for electrons. It uncovers diffuse scattering which can originate from protein disorder or dynamics, providing additional information that improves accurate modeling and refinement^17^. As energy filtering removes a major obstacle to better data quality, other sources of noise that remain can be addressed more effectively and potentially lead to further improved accuracy in macromolecular electron crystallography.

## Data availability

Coordinates and structure factors have been deposited to the PDB.

## Acknowledgements

This study was supported by the National Institutes of Health P41GM136508 and the Department of Defense HDTRA1-21-1-0004 and the Howard Hughes Medical Institute.

## Methods

### Crystallization

Proteinase K powder from *Engyodontium/tritirachium album* was purchased from Sigma Aldrich and used without further purification. Crystals were grown in batch by dissolving 40 mg/ml proteinase K in 20 mM MES-NaOH pH 6.5. The protein solution was mixed at a 1:1 ratio with a precipitant solution of 0.5 M NaNO_3_, 0.1 M CaCl_2_, 0.1 M MES-NaOH pH 6.5. The mixture was incubated at 4 °C. Microcrystals with dimensions ranging between 8-12 μm grew within 24h.

### Sample preparation

Standard holey carbon electron microscopy grids (Quantifoil, Cu 200 mesh, R2/2) were glow discharged for 30 s at 15 mA on the negative setting. Samples were prepared using a Leica GP2 vitrification device set at 4 °C and 90% humidity. For each sample, 3 μl of crystal solution was deposited onto the grid, incubated for 10s, and any excess liquid was blotted away from the back side. Next, the sample was soaked for 30s with 3 μl cryoprotectant solution containing 30% glycerol, 250 mM NaNO_3_, 50 mM CaCl_2_, 60 mM MES-NaOH pH 6.5. After incubation, any excess solution was blotted away using filter paper for a second time. Immediately after, the grid was rapidly vitrified by plunging it into liquid ethane. Grids were stored in a liquid nitrogen dewar prior to further use.

### Focused ion beam milling

Grids were loaded onto a Helios Hydra 5 CX dual-beam plasma FIB/SEM (Thermo Fisher Scientific). Prior to milling, grids were coated with a thin protective layer of platinum for 45s using the gas injection system. Microcrystals of proteinase K were machined using a stepwise protocol to an optimal thickness of approximately 300 nm using a 30 kV Argon plasma ion beam. Coarse milling steps were performed using a 2.0 nA current to a thickness of approximately 3 μm. Finer milling steps at 0.2 nA were used to thin the lamellae to 600 nm. Final polishing steps were performed at 60 pA down to 300 nm thickness. Grids were cryo-transferred immediately after to the TEM for data collection. Grids were rotated by 90° relative to the milling direction such that the rotation axis on the microscope is perpendicular to the milling direction.

### Hardware setup

Data were collected on a Titan Krios G3i TEM (Thermo Fisher Scientific) equipped with a X-FEG operated at an acceleration voltage of 300 kV, a post-column Selectris energy filter, and a Falcon 4i direct electron detector. The microscope was aligned for low flux density conditions using the 50 μm C2 aperture, spot size 11, and gun lens setting 8 for a less bright but more coherent illumination. A parallel electron beam of 10 μm diameter was used for data collection. The flux density at these conditions is approximately 0.002 e^-^/Å^2^/s. The energy spread of the emitted electrons was characterized at *ΔE* = 0.834 eV +/-0.006 eV at FWHM. The zero loss peak of the energy filter was first centered in imaging mode. On our system, there is an offset in the position of the zero-loss peak when switching between imaging and diffraction mode. Therefore, the energy filter slit was carefully offset manually in defocused diffraction mode to align it with the selected area (SA) aperture used for diffraction data collection.

### Data collection

Electron-counted and energy-filtered MicroED data were collected using the continuous rotation method. Diffraction data were collected using a 150 μm SA aperture, corresponding to a beam diameter of ∼3 μm at the sample plane as defined by the aperture. The energy filter zero-loss peak was centered in defocused diffraction at the 5 and 10 eV slit settings. The effective sample to detector distance was calibrated at 1402 mm using a standard evaporated aluminum grid (Ted Pella). Crystals were rotated at a slow angular increment of 0.0476 ∘/s covering a total tilt range of 20.0∘. Data were collected over 420 s exposures at a total fluence of ∼0.84 e^-^/Å^2^. Diffraction stills were taken using the same settings without any stage rotation from stationary crystals. No beam stop post energy filter was used. Even without a beam stop, the low resolution reflections close to the center, where most of the inelastic scattering normally accumulates, are in sharp contrast to the low background. Data were recorded on a Falcon 4i direct electron detector in electron counting mode operating at an internal frame rate of 320 Hz. The proactive dose protector was manually disabled. Raw data were written in electron event representation (EER) format with an effective readout speed of 308 frames per second.

### Data processing

Individual MicroED datasets in EER format were binned by two and converted to SMV format using the MicroED tools (available at https://cryoem.ucla.edu), after summing batches of 308 frames and applying post-counting gain corrections. Individual MicroED datasets were processed using XDS^18^. Data were integrated up to a cross correlation between two random half sets that was still significant at the 0.1% level^19^. Individual datasets were analyzed and merged using XSCALE^18^ and XSCALE_ISOCLUSTER^20^. The merged data were truncated at a mean *I/σI* ≥ 1.0 and a CC_1/2_ of 33.0 % in the highest resolution shell. Data were merged using Aimless^21^.

### Structure solution and refinement

The structure was phased by molecular replacement using electron scattering factors in Phaser^22^. The model was inspected and built using Coot^23^. Two calcium ions and one nitrate were placed, and a total of 21 residues were modelled in alternate conformations. The structure was refined using REFMAC5^24^ and phenix.refine^25^. Refinement was done using electron scattering factors and individual anisotropic *B*-factors for all atoms except hydrogens. Hydrogen omit maps were calculated using REFMAC5. Difference peaks higher than 2σ and 3σ in the omit map within less than 1.0 Å from any known riding positions were identified as potential hydrogen atoms.

## Figures and Tables

**Extended Data Figure 1.**
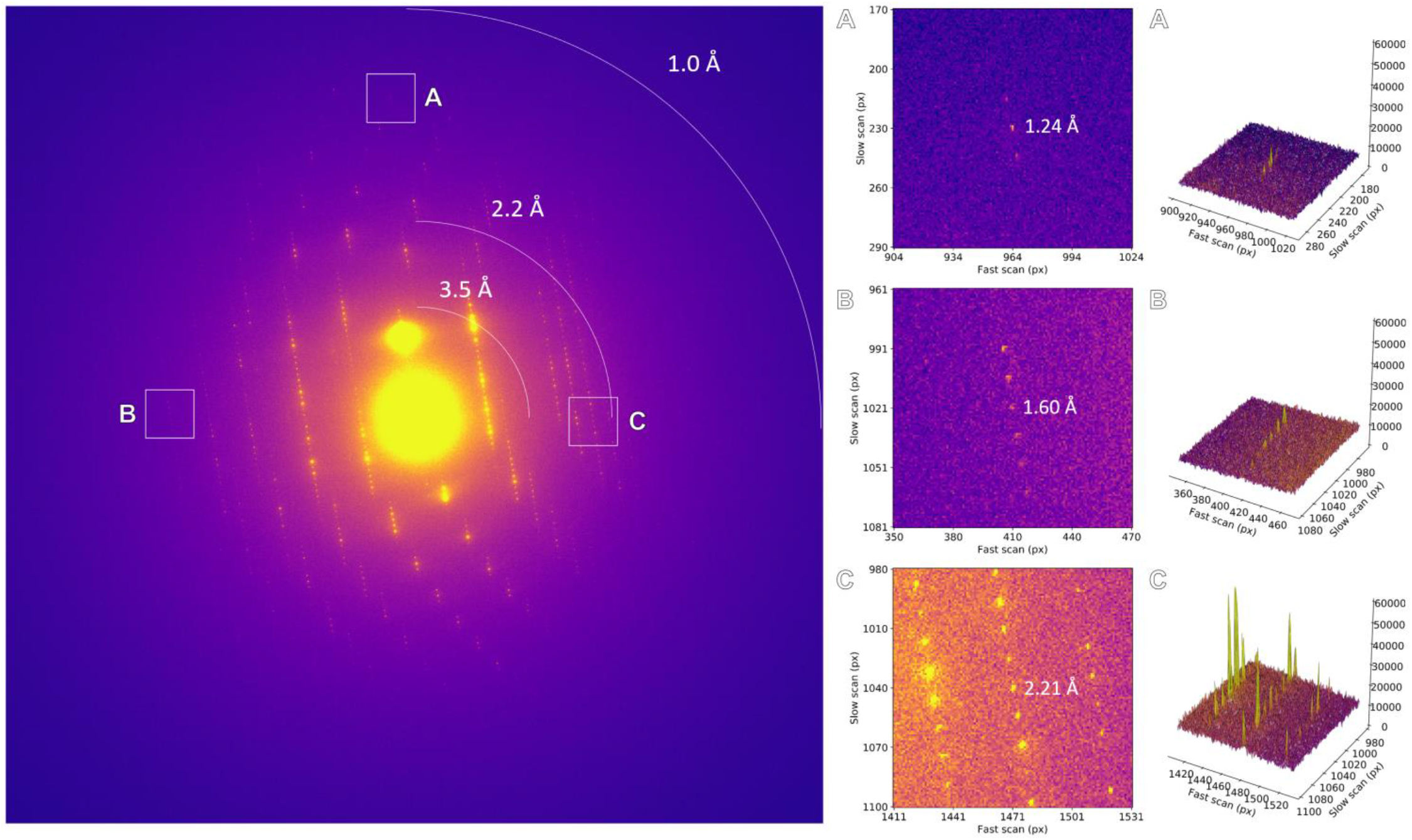
Unfiltered diffraction pattern of a stationary proteinase K lamella. Areas highlighted by the insets are magnified and the corresponding peak profiles are plotted.

**Extended Data Figure 2.**
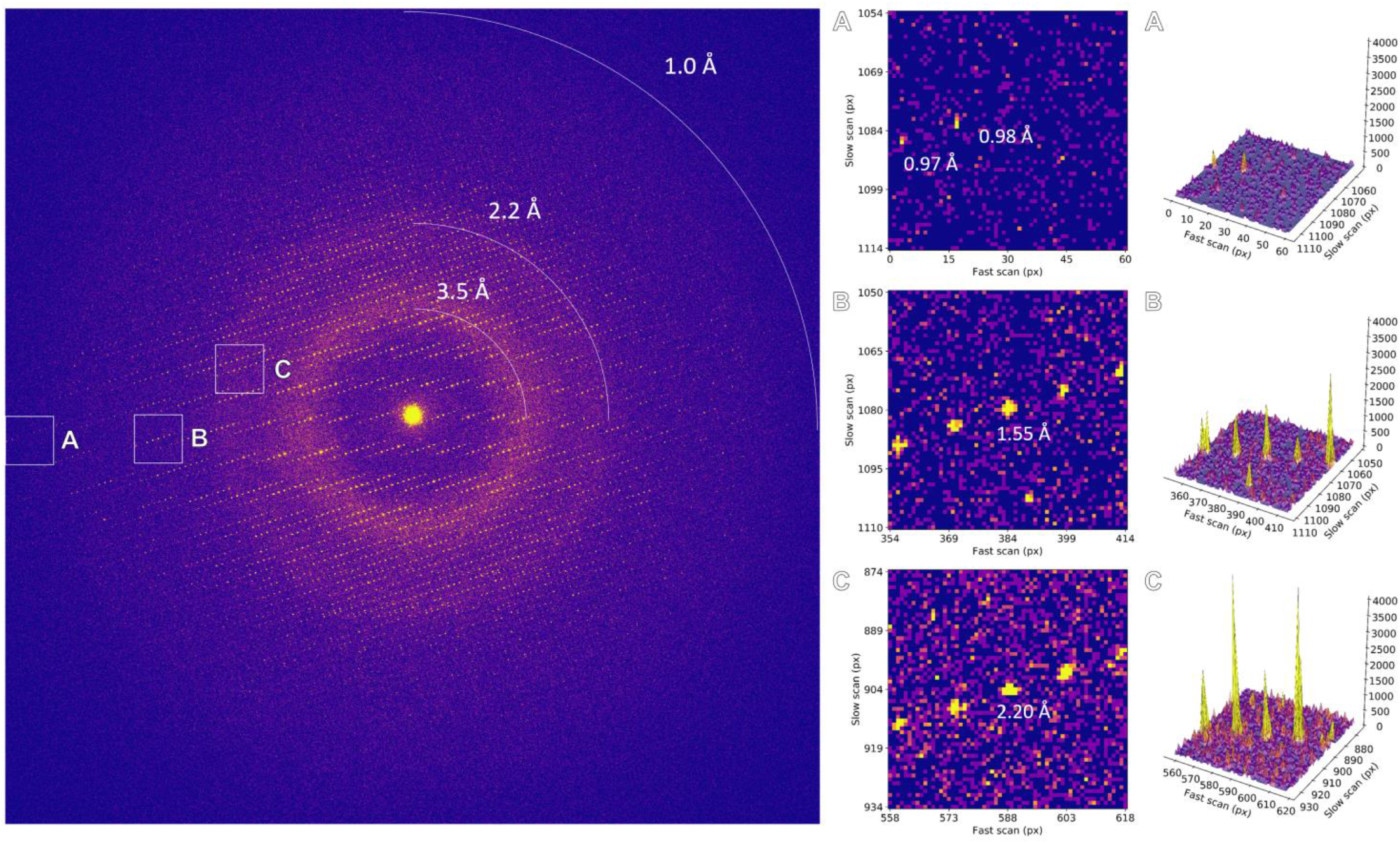
Individual frame from MicroED data collection using a 10 eV energy filter slit setting. Areas highlighted by the insets are magnified and the corresponding peak profiles are plotted.

**Extended Data Figure 3.**
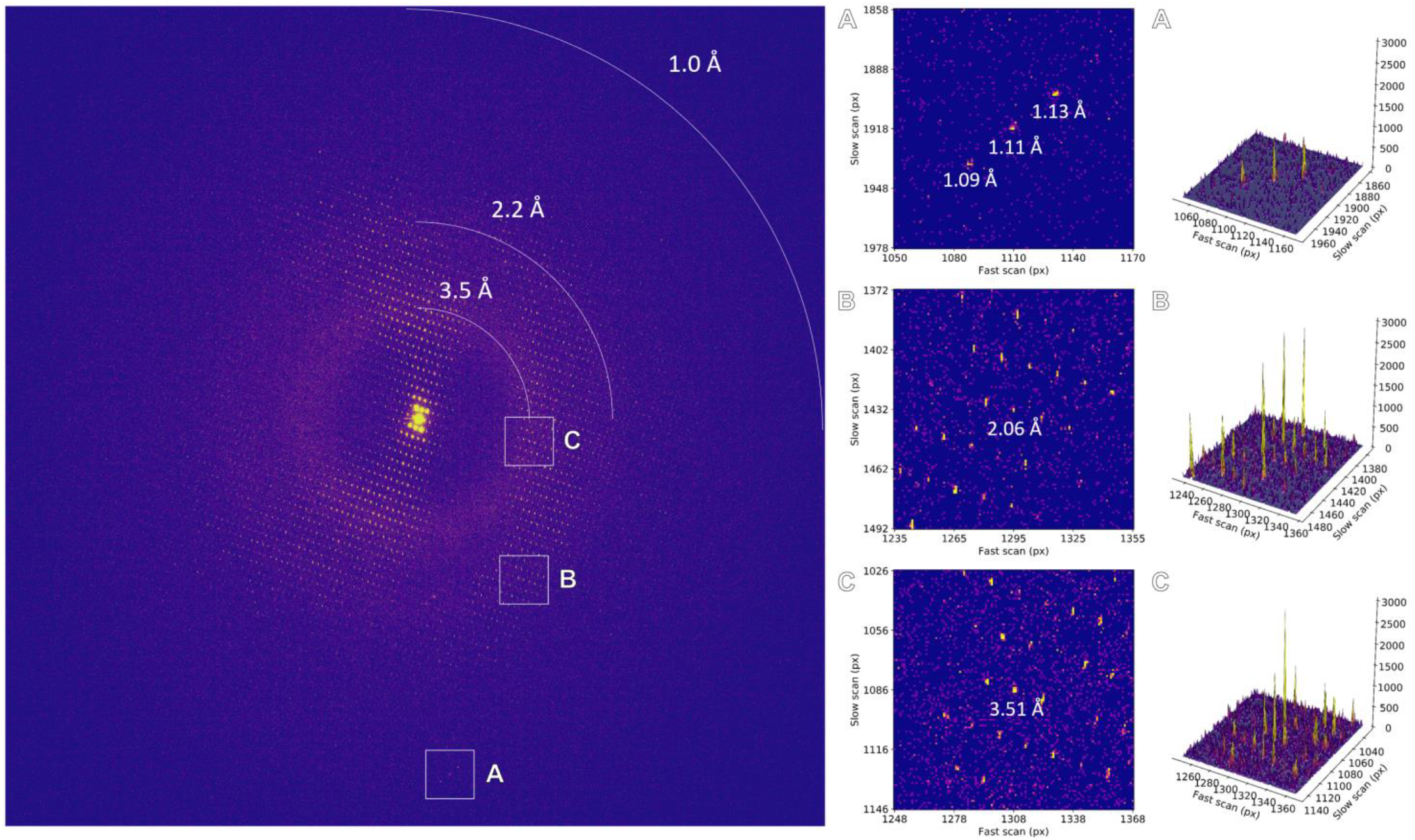
Individual frame from MicroED data collection using a 5 eV energy filter slit setting. Areas highlighted by the insets are magnified and the corresponding peak profiles are plotted.

**Extended Data Figure 4.**
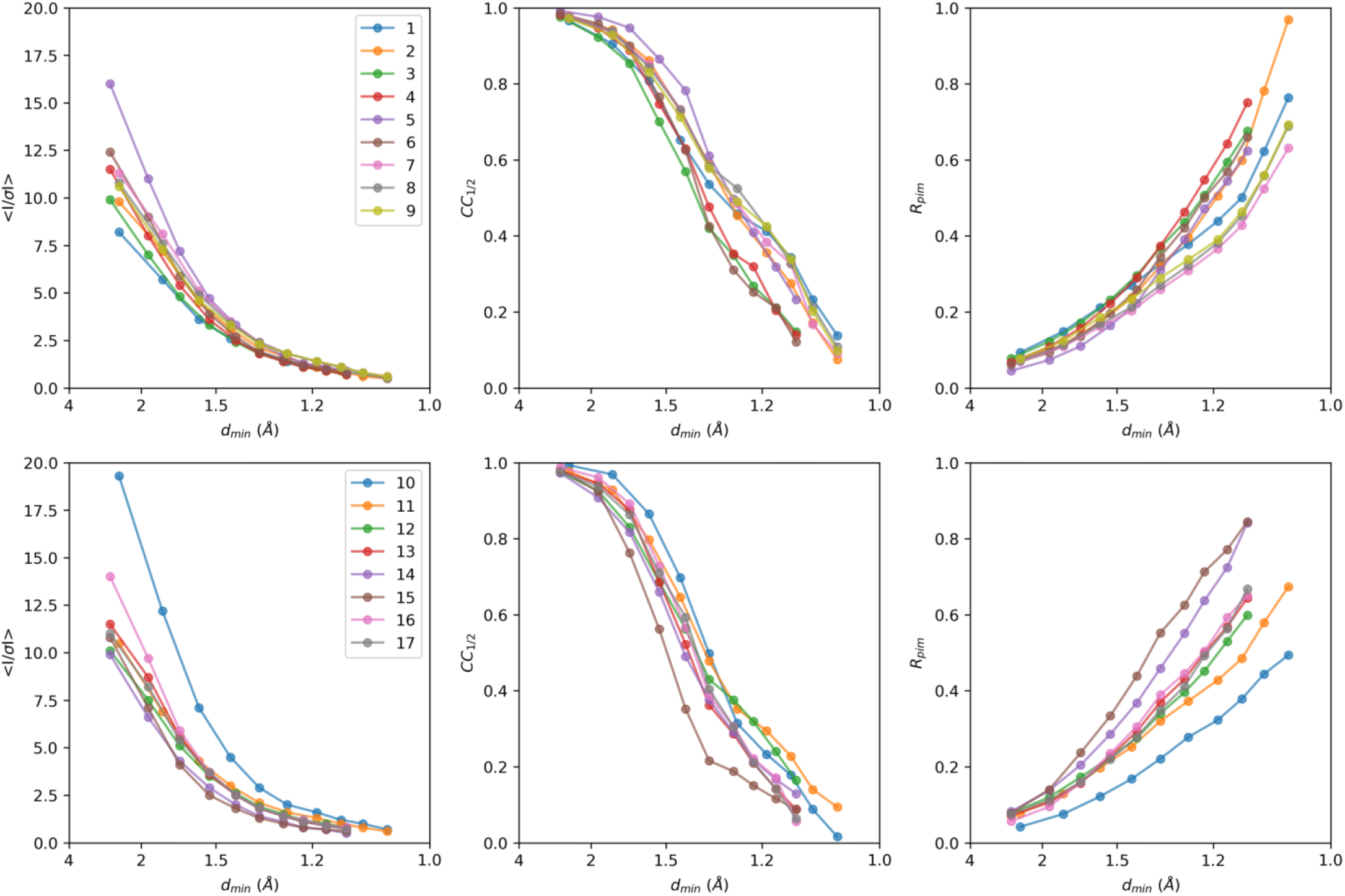
Intensity statistics for individual MicroED datasets, featuring from left to right *I/σI*, CC_1/2_, and Rpim as function of the resolution.

**Extended Data Table 1.** Data processing and refinement statistics.

| **Data collection** |  |
| --- | --- |
| No. of crystals | 17 |
| Space group | $P4_32_12$ |
| Unit cell dimensions |  |
| $a, b, c$ (Å) | 66.92, 66.92, 107.56 |
| $\alpha, \beta, \gamma$ (°) | 90, 90, 90 |
| Resolution (Å)* | 56.90-1.09 (1.12-1.09) |
| Observed reflections | 2,811,895 (95,185) |
| Unique reflections | 98,530 (6,664) |
| Multiplicity | 28.5 (14.3) |
| Completeness (%) | 97.5 (90.3) |
| $R_{\text{merge}}$ | 0.284 (1.154) |
| $\langle I/\sigma I \rangle$ | 7.8 (1.4) |
| $CC_{1/2}$ (%) | 99.5 (33.0) |
| **Refinement** |  |
| Resolution (Å) | 56.82-1.09 |
| No. of reflections | 98,228 |
| $R_{\text{work}}/R_{\text{free}}$ | 0.149/0.182 |
| No. of atoms |  |
| Protein | 2193 |
| Ligand/ions | 6 |
| Waters | 357 |
| Hydrogen omit maps |  |
| No. of hydrogen atoms | 1942 |
| Peaks at $\geq 3\sigma$ | 737 (37.9%) |
| Peaks at $\geq 2\sigma$ | 1280 (69.7%) |
| $\langle B \rangle$ factors (Å<sup>2</sup>) | |
| Overall | 11.76 |
| Protein | 9.17 |
| Ligand/ions | 11.67 |
| Waters | 27.67 |
| R.m.s. deviations |  |
| Bond lengths (Å) | 0.016 |
| Bond angles (°) | 1.24 |

\*Values in parenthesis are for the highest resolution shell. Data were truncated at a mean $I/\sigma I \geq 1.0$ and a cross correlation between two random half sets that is statistically significant at $CC_{1/2} \sim 33\%$ .

## Notes

### Competing Interest Statement

The authors have declared no competing interest.

